# Blocking the alternative sigma factor RpoN reduces virulence of *Pseudomonas aeruginosa* isolated from cystic fibrosis patients and increases antibiotic sensitivity in a laboratory strain

**DOI:** 10.1101/340307

**Authors:** MG Lloyd, JL Vossler, CT Nomura, JF Moffat

## Abstract

Multidrug-resistant organisms (MDROs) are increasing in the health care setting, and there are few antimicrobial agents available to treat infections caused by these bacteria. *Pseudomonas aeruginosa* is an opportunistic pathogen in burn patients and individuals with cystic fibrosis (CF), and a leading cause of nosocomial infections. *P. aeruginosa* is inherently resistant to many antibiotics and can develop or acquire resistance to others, limiting options for treatment. *P. aeruginosa* has virulence factors that are regulated by sigma factors in response to the tissue microenvironment. The alternative sigma factor, RpoN (σ^54^), regulates many virulence genes and is linked to antibiotic resistance. Recently, we described a cis-acting peptide, RpoN*, which acts as a “molecular roadblock”, binding RpoN consensus promoters at the −24 site and blocking transcription. RpoN* reduces virulence of *P. aeruginosa* laboratory strains both *in vitro* and *in vivo,* but its effects in clinical isolates was not known. We investigated the effects of RpoN* on phenotypically varied *P. aeruginosa* strains isolated from cystic fibrosis patients. RpoN* expression reduced motility, biofilm formation, and pathogenesis in a *P. aeruginosa – C. elegans* infection model. RpoN* expression increased susceptibility to several beta-lactam based antibiotics in the lab strain *P. aeruginosa* PA19660 *Xen5*. Here, we show that using a cis-acting peptide to block RpoN consensus promoters has potential clinical implications in reducing virulence and enhancing the activity of antibiotics.

## Introduction

Multidrug-resistant organisms (MDROs) are an increasing problem in the healthcare setting. Both Gram-negative and Gram-positive MDROs are prevalent globally (1, 2). There are few or no antimicrobial agents available for treatment of infections caused by these bacteria (3). *Pseudomonas aeruginosa*, a Gram-negative, opportunistic pathogen is a leading cause of nosocomial infections and is associated with infections in burn patients (4, 5). *P. aeruginosa* is also responsible for colonizing the respiratory tract and causing chronic infections in individuals with cystic fibrosis (CF) (6). It is the most common pathogen isolated from individuals with CF, and is a major source of morbidity and mortality (7–10).

In CF patients, *P. aeruginosa* undergoes a transformation from a non-mucoid form upon initial colonization of the lungs to a mucoid form as the disease progresses. This results in a chronic debilitating pulmonary infection characterized by the overexpression of alginate. Mucoid strains synthesize large quantities of alginate exopolysaccharide, enhancing biofilm formation and protecting *P. aeruginosa* from antibiotics or the immune response (11), possibly through formation of microcolonies (12, 13). While aggressive prevention regimens have led to a decline in prevalence of *P. aeruginosa* in CF patients, multidrug resistant strains are still prevalent and occurred in 19.4% of CF infections in 2015 (14). *P. aeruginosa* is inherently resistant to a number of antibiotics (15, 16). It can also acquire resistance through exogenous resistance genes via horizontal gene transfer or mutations (17), limiting available treatment options. Antimicrobial development is directed toward alternative treatments and novel targets. Promising strategies include enhancing the activity of currently available antibiotics and decreasing virulence of the bacteria once an infection occurs (18–25).

*P. aeruginosa* virulence is caused by many factors, including production of toxins, proteases, phospholipases, the presence of pili and flagella, and biofilm formation (26). This virulence is regulated by a network of transcription factors, such as sigma factors RpoS and RpoN, and quorum sensing regulators (27). The alternative sigma factor, σ^54^ or RpoN, regulates nitrogen assimilation, quorum sensing, motility, and biofilm formation (28–33). RpoN regulation was recently linked to *P. aeruginosa* tolerance to several antibiotics (34–36). RpoN binds to specific promoters with conserved −24, −12 sequences upstream of RpoN-regulated genes throughout the genome and is a key virulence regulator (37). The specific and conserved nature through which RpoN controls its regulon led us to develop the RpoN molecular roadblock, RpoN*. RpoN* is a cis-acting peptide that specifically binds the −24 site of RpoN consensus promoters, blocking transcription by RpoN and other factors (38). When RpoN* is expressed in *P. aeruginosa* laboratory strains, transcription is affected globally and virulence is attenuated (38). RpoN* also affects virulence in an RpoN deletion strain of *P. aeruginosa* PAO1, demonstrating its ability to attenuate gene expression by repressing expression of genes located downstream of RpoN promoters (38). This strategy of blocking multiple promoters throughout the *P. aeruginosa* genome may be an effective method to combat virulence and evade development of resistance.

*P. aeruginosa* isolated from CF patients are phenotypically and genetically varied (39, 40). Many *P. aeruginosa* clinical isolates have mutations, including deletion or loss of function, in the *rpoN* gene (41, 42). It was not known how the cis-acting RpoN* peptide would affect virulence phenotypes in *P. aeruginosa* clinical isolates, particularly in strains that do not express or have low levels of RpoN. In this study, we describe the effects of RpoN* on *in vitro* and *in vivo* virulence of *P. aeruginosa* isolated from CF patients and its effects on antibiotic resistance. Expression of RpoN* reduced virulence-associated phenotypes in clinical isolates and improved *P. aeruginosa* susceptibility to multiple antibiotics. This study demonstrates that RpoN* has potential clinical applications and potentially represents an effective strategy to combat both antibiotic resistance and infections with *P. aeruginosa* in CF patients.

## Results

### Virulence phenotypes were variable in P. aeruginosa isolates from CF patients

*P. aeruginosa* isolated from different CF patients or within the same CF patient have varied phenotypes and genotypes (39, 40). *P. aeruginosa* adapts over time, leading to mutations and changes in expression of genes related to motility, quorum sensing, and overall virulence (41, 43). To determine the virulence-related phenotypic profiles of the strains used in this study (Table 1), each *P. aeruginosa* patient isolate was evaluated for motility and biofilm formation, compared to the positive, virulent control strain *P. aeruginosa* PA19660 *Xen5*. Several patient isolates were highly motile in the swimming assay (flagella), including SCH0057-7, SCH0256-1, SCH0354-1 and UUH0201, while others were nonmotile (Fig 1A). Most strains were motile in the twitching assay (pili) and produced moderate biofilms, with SCH0254-118 migrating the furthest (Fig 1B) and forming the most extensive biofilm (Fig 1C). SCH0254-116, SCH0397-3, and UUH0202 did not form biofilms.

**Table 1.** *P. aeruginosa* strains used in this study.

| Location | Strain | Source | CF mutations | Reference |
| --- | --- | --- | --- | --- |
| <u>Laboratory</u> |  |  |  |  |
|  | PAO1-M | --- | n/a | C. Manoil (27) |
|  | PAO1-S | --- | n/a | D. Haas (15) |
| | $\Delta rpoN$ | --- | n/a | D. Haas (15) |
|  | PA19660 Xen5 | Septicemia | n/a | Perkin Elmer |
| <u>Clinical Isolate</u> |  |  |  |  |
| *Seattle Children's Hospital (SCH)* |  |  |  |  |
| | SCH0057-7 | unknown | $\Delta F508$ / $\Delta F508$ | This study |
| | SCH0254-23 | unknown | $\Delta F508$ / unknown | This study |
| | SCH0254-116 | unknown | $\Delta F508$ / unknown | This study |
| | SCH0254-118 | unknown | $\Delta F508$ / unknown | This study |
| | SCH0256-1 | sputum | $\Delta F508$ / $\Delta F508$ | This study |
|  | SCH0338-38 | sputum | unknown / unknown | This study |
| | SCH0354-1 | sputum | $\Delta F508$ / G551D | This study |
| | SCH0397-3 | unknown | $\Delta F508$ / unknown | This study |
| | SCH03269 | sputum | $\Delta F508$ / $\Delta F508$ | This study |
| *Upstate University Hospital (UUH)* |  |  |  |  |
| | UUH0101 | sputum | $\Delta F508$ / unknown | This study |
| | UUH0201 | sputum | $\Delta F508$ / $\Delta F508$ | This study |
| | UUH0202 | sputum | $\Delta F508$ / $\Delta F508$ | This study |

**Figure 1.**
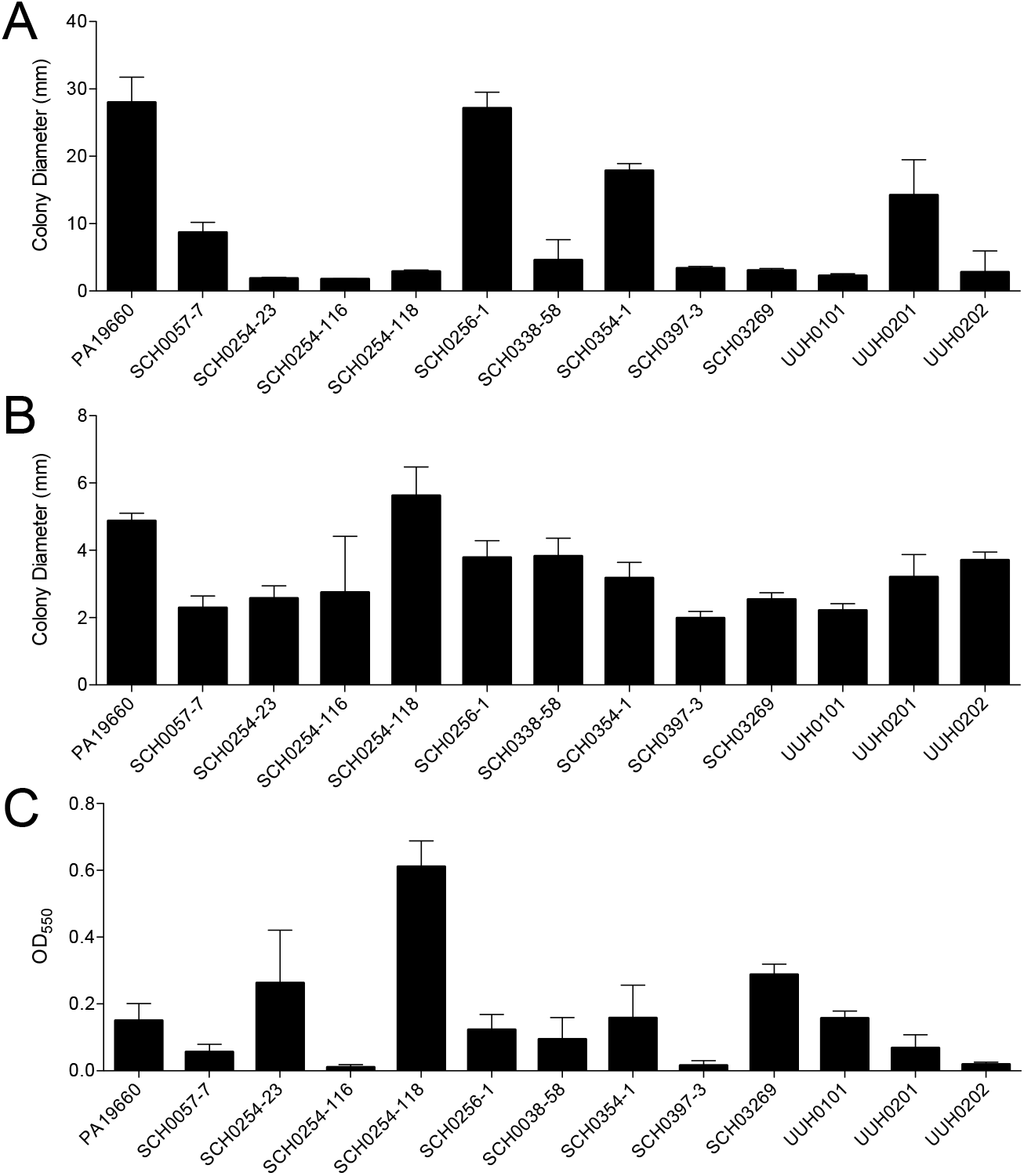
Characterization of virulence phenotypes of *P. aeruginosa* strains isolated from cystic fibrosis patients. *P. aeruginosa* CF patient isolates were compared to laboratory strain PA19660 *Xen5*. All assays were conducted at 37°C for 24 h. (A) Colony diameter of swimming, or flagellar, motility assay conducted on soft (0.3%) agar. (B) Twitching, or pili, motility assay conducted on semi-hard (1.3%) agar. Colony diameter was measured across point of inoculation to the edges of bacteria colony. (C) Biofilm formation assay was conducted in 96-well microtiter plates. Biofilms were stained with crystal violet (0.1%), solubilized in ethanol (95%), and absorbance measured at OD550. Bars are the mean ± SD; n = 5 to 6 replicates in motility assays and n = 10 in biofilm assay. Each assay was performed at least three separate times and representative results are shown.

The pathogenesis of patient isolates was evaluated in a *P. aeruginosa – C. elegans* infection model. All patient isolates were compared to *E. coli* OP50, an avirulent negative control. SCH0057-7 was the most pathogenic in the paralytic killing assay, which is mediated by hydrogen cyanide production (44, 45) (Fig 2A). Other strains were moderately pathogenic, including SCH0256-1, SCH0354-1, SCH0397-3, and UUH0202. SCH0057-7, SCH0338-58, and UUH0202 were highly pathogenic in the slow killing assay, which mimics establishment and proliferation of an infection and is mediated by the *lasR, gacA, lemA,* and *ptsP* genes (46), while UUH0201 was moderately pathogenic (Fig 2B). As expected, the virulence-associated phenotypes of patient isolates varied widely *in vitro* and *in vivo*.

**Figure 2.**
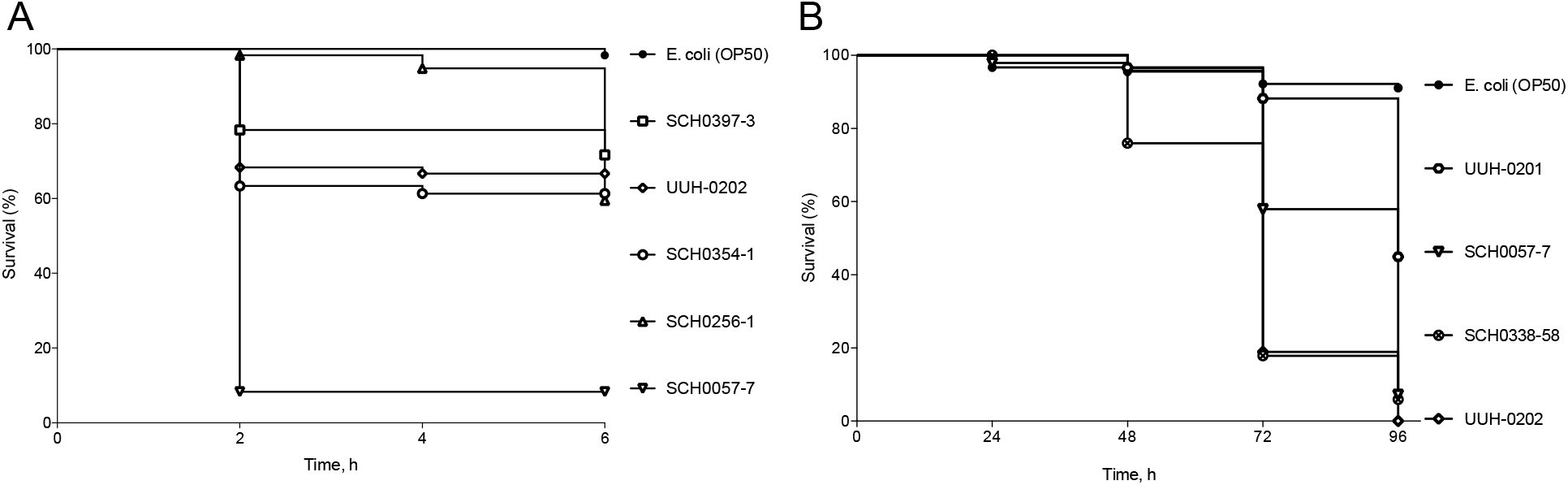
Pathogenesis of *P. aeruginosa* isolated from cystic fibrosis patients in *P. aeruginosa – C. elegans* infection model. Kaplan-Meier survival curves for *P. aeruginosa – C. elegans* infection assays. (A) Paralytic killing assay on BHI agar. Assay was conducted at room temperature and worm status scored every 2 h. (B) Slow killing assay on modified NGM agar (0.35% bactopeptone, 2% bactoagar). Assay was conducted at 20°C and worm status scored every 24 h. Strains used included CF patient isolates, and *E. coli* for reference. n = 48 to 90 worms per strain.

### RpoN protein levels varied among patient isolates

Others reported that the *rpoN* gene was mutated or lost in approximately 20% of *P. aeruginosa* isolates from CF patients (41). Loss or mutation in the *rpoN* gene can result in phenotypes similar to those observed in the patient isolates evaluated here (29, 31, 32). Thus, we evaluated relative protein levels of RpoN in these patient isolates by western blot. RpoN levels were moderately high in the positive control *P. aeruginosa* PAO1-S, while low or minimal protein levels were detected in the isogenic Δ*rpoN* mutant negative control (Fig 3). The low level of background in the Δ*rpoN* mutant is likely due to nonspecific antibody binding. RpoN levels varied in the CF patient isolates, with high levels in SCH0057-7, SCH0397-3, and UUH0201; intermediate levels in SCH0254-116, SCH0338-58, and UUH0202; and low levels in SCH0254-23, SCH0254-118, SCH0256-1, SCH0354-1, SCH03269, and UUH0101.

**Figure 3.**
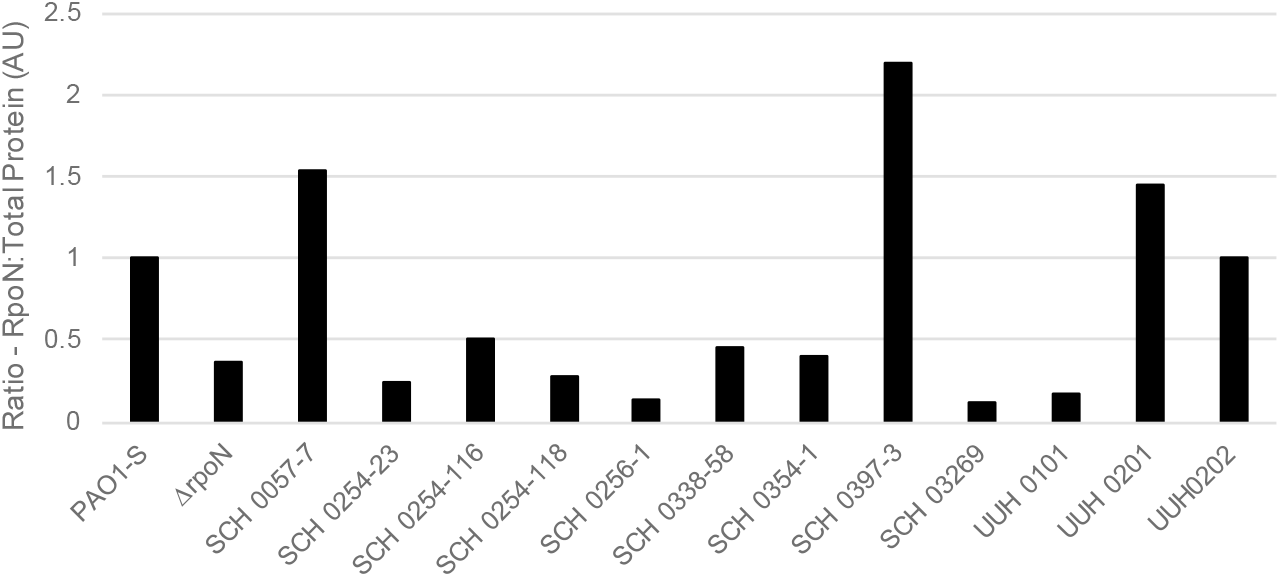
RpoN protein in *P. aeruginosa* isolates is highly varied. Immunoblot analysis of RpoN expression in *P. aeruginosa* CF patient isolates and laboratory strains PAO1-S and Δ*rpoN*. Immunoblots performed on 10% Mini-PROTEAN^®^ TGX Stain-Free™ gels (BioRad). RpoN protein levels were calculated by comparing measured total protein in each lane to the measured RpoN band (presented as arbitrary units (AU)). Values were normalized against PAO1-S. This graph is representative of immunoblots from multiple bacterial cultures and western blot analyses.

### RpoN* expressed in CF patient isolates reduced virulence-associated phenotypes in vitro

The effect of RpoN* expression on motility and biofilm formation in patient isolates was not known. Unfortunately, some patient isolates could not be transformed, and so only four isolates were evaluated for the effects of RpoN* expressed from a plasmid. SCH0057-7, SCH0256-1, SCH0338-58, and SCH0354-1 were transformed with a plasmid expressing RpoN* or the empty vector and selected with gentamicin. If RpoN* affected transcription of virulence-related genes in different genetic backgrounds as previously reported (38), we expected attenuation of virulence-related phenotypes in *P. aeruginosa* CF patient isolates. RpoN* significantly reduced colony diameter in all four patient isolates in the swimming motility assay (Student’s t-test, ** p≤0.01, ***p≤0.0001) (Fig 4A). RpoN* significantly reduced colony diameter in SCH0057-7, SCH0256-1, and SCH0338-58 in the twitching motility assay (Student’s t-test, ** p≤0.01, ***p≤0.0001) (Fig 4B). Colony diameter varied widely in SCH0354-1 when RpoN* was expressed and was always smaller than with empty vector, although the difference was not significant. In the biofilm formation assay, RpoN* significantly reduced biofilm formation by SCH0057-7 and SCH0256-1 (Student’s t-test, p≤0.0001) (Fig 4C). Thus RpoN* reduced virulence-associated phenotypes of *P. aeruginosa* isolated from CF patients.

**Figure 4.**
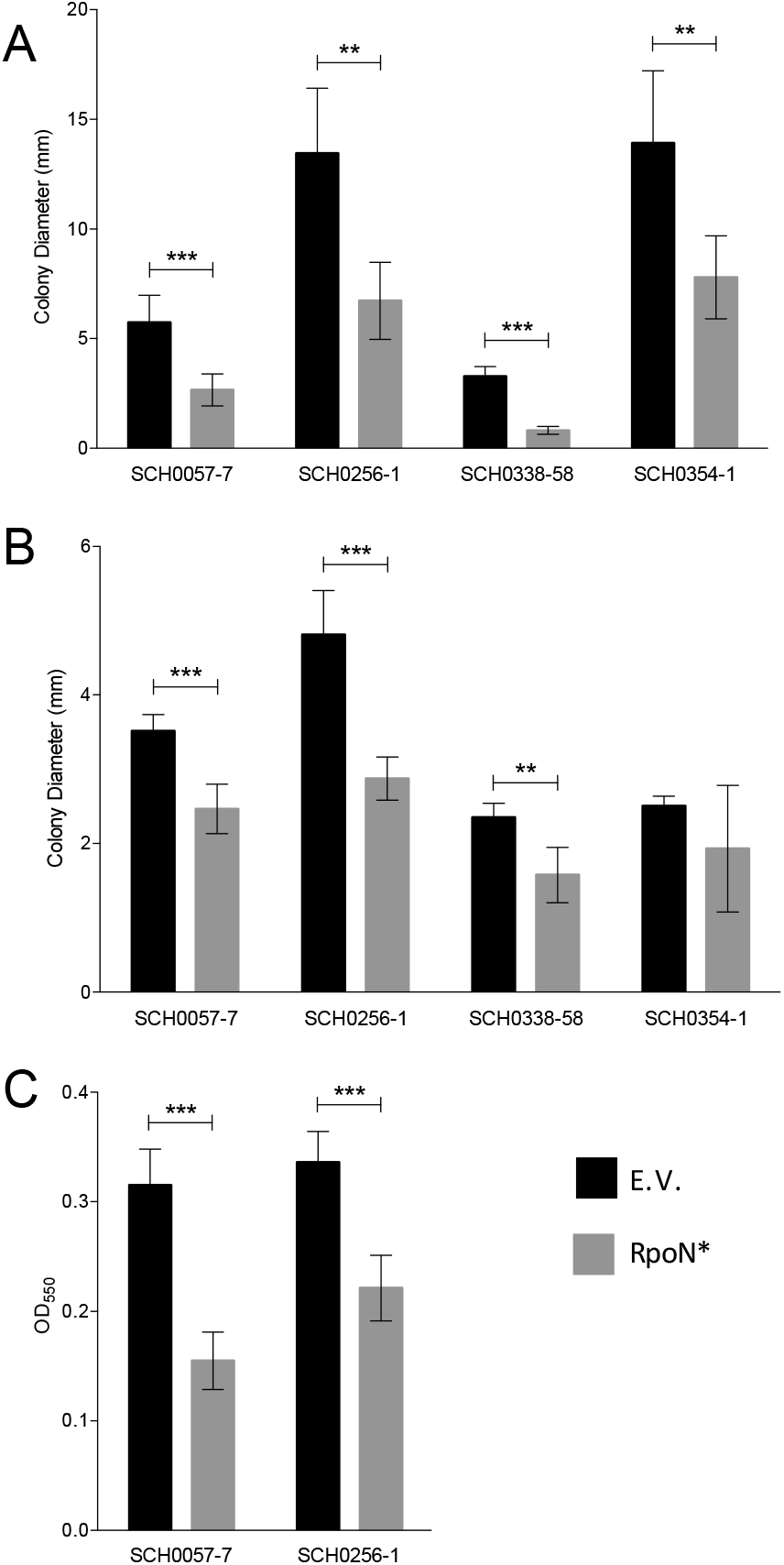
RpoN* expression decreases motility and biofilm formation. *P. aeruginosa* CF patient isolates with empty vector (E.V., black bars), or with RpoN*-expression vector (gray bars). Media was supplemented with gentamicin (30 mg/L), and IPTG (1 mM) when applicable, and all assays were conducted at 37°C for 24 h. (A) Colony diameter of swimming, or flagellar, motility assay conducted on soft (0.3%) agar. (B) Colony diameter of twitching, or pili, motility assay conducted on semi-hard (1.3%) agar. (C) Biofilm formation assay conducted in 96-well microtiter plates. Bars are the mean ± SD (Student’s t-test, *** p ≤ 0.0001; ** p ≤ 0.01). n = 4 to 5 replicates in motility assays; n = 12 in biofilm assays.

### RpoN* expression increased worm survival in P. aeruginosa – C. elegans infection model

Initial evaluation of patient isolates revealed a single *P. aeruginosa* strain, SCH0057-7, that was both transformable and pathogenic in the *P. aeruginosa – C. elegans* infection assay. Therefore, effects of RpoN* on pathogenesis of SCH0057-7 were evaluated using the paralytic killing assay, which is based on *P. aeruginosa* hydrogen cyanide production and mimics conditions in the CF lung (44, 45). Wild-type *P. aeruginosa* SCH0057-7 was the positive, virulent control and *E. coli* was the negative, avirulent control. The test conditions were *P. aeruginosa* SCH0057-7 expressing RpoN* or carrying the empty vector plasmid. If RpoN* affected virulence-related phenotypes in *P. aeruginosa* SCH0057-7, then we expected increased survival of *C. elegans*. Wild type SCH0057-7 and with the empty vector killed approximately 80% of *C. elegans* (Fig 5). In contrast, RpoN* expression significantly increased *C. elegans* survival (Mantel-Cox Log-Rank Test, p≤0.0001). Thus, RpoN* expression reduced pathogenesis of a patient isolate in a *P. aeruginosa – C. elegans* infection model.

**Figure 5.**
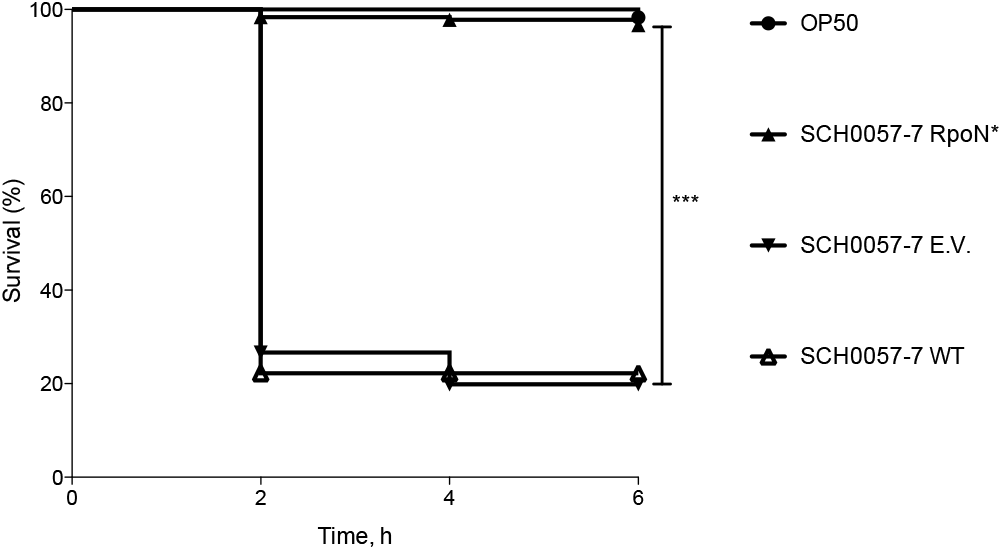
RpoN* promotes *C. elegans* survival in paralytic killing assay. *P. aeruginosa* CF isolate SCH0057-7 wild type, with empty vector (E.V.), or with RpoN*-expression vector, were compared to avirulent *E. coli* OP50. Paralytic killing assay was conducted on BHI agar supplemented with gentamicin (30 mg/L) and IPTG (1 mM), when applicable. Assay was conducted at room temperature, and worm status scored every 2 h. Kaplan-Meier survival curves represent combined survival of three separate assays (exception: *E. coli)*. Mantel-Cox log-rank test was used to analyze E.V. and RpoN* curves (*** p≤0.0001). n = 180 worms per SCH0057-7 condition, n = 60 worms for *E. coli*.

### RpoN* increased antibiotic susceptibility in vitro

Antibiotic resistance is a problem in CF patients with *P. aeruginosa* infections (47–49). We previously reported that RpoN* alters transcription of several genes involved in multidrug efflux pumps that confer natural resistance (38). Additionally, RpoN is implicated in tolerance to various classes of antibiotics (34–36). We evaluated the effects of RpoN* on antibiotic susceptibility using a MicroScan Neg MIC 43 panel. The test conditions were *P. aeruginosa* PA19660 *Xen5* that was mock-transformed, or transformed with the empty vector or RpoN* plasmid. We expected that RpoN* would improve antibiotic susceptibility of *P. aeruginosa*. In PA19660 Xen5 mock-transformed or with the empty vector, antibiotic susceptibility profiles were the same, except for gentamicin, which increased in the empty vector strain due to the GM^R^ selection marker (data not shown). In PA19660 *Xen5* expressing RpoN*, susceptibility to five beta-lactam antibiotics was improved 2-to 4-fold (Fig 6). These were cefotaxime, cefepime, and ceftazidime (three cephalosporins), piperacillin (a ureidopenicillin), and imipenem (a carbapenem). Susceptibility to some antibiotics was unchanged (data not shown). The results demonstrate that RpoN* expression increased *P. aeruginosa* susceptibility to several antibiotics.

**Figure 6.**
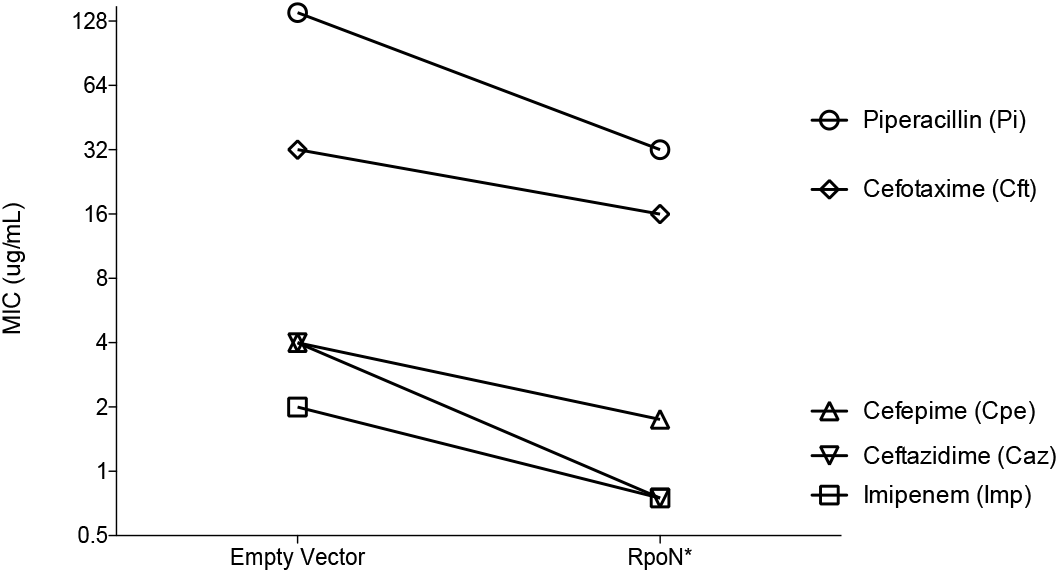
RpoN* expression increases susceptibility to antibiotics. Overnight cultures of *P. aeruginosa* PA19660 Xen5 with empty vector or expressing RpoN*, were diluted in fresh LB broth, added to wells of MicroScan Neg MIC 43 panel, and incubated at 35°C for 16-20 h. Media was supplemented with gentamicin (30 mg/L) and IPTG (1 mM). Each point is the average of two biological replicates per experiment and the data shown are representative of three separate experiments.

## Discussion

Here, we confirm and expand results of previous studies by showing the ability of RpoN* to abrogate virulence phenotypes in *P. aeruginosa* isolates from CF patients and to improve susceptibility to several antibiotics. Our working model of the mechanism of action of RpoN* is that it binds the −24 promoter consensus sites, blocking transactivation by RpoN and other sigma factors. By altering the transcriptome, RpoN* reduced virulence in well-characterized laboratory strains (38). Thus, the motivation for this study was to understand the clinical relevance of RpoN*. We demonstrated that RpoN* expressed in CF patient isolates reduced motility and biofilm formation *in vitro,* independently of RpoN protein levels. The RpoN* molecular roadblock protected *C. elegans* from a highly virulent *P. aeruginosa* patient isolate in an *in vivo* infection model. RpoN* also improved *P. aeruginosa* susceptibility to antibiotics.

*P. aeruginosa* isolated from CF patients are highly variable (39, 40), with the *rpoN* gene often mutated or lost (41). The patient isolates evaluated in this study had a broad range of motility, biofilm formation, RpoN protein levels, and virulence in *C. elegans*. There was no correspondence between most *in vitro* phenotypes, *in vivo* pathogenesis, and RpoN levels (Fig. S1). The only correlation observed was between twitching or pili-associated motility and biofilm formation (Supplemental Fig 1F, p=0.0357, R^2^=0.37050). Other studies suggested that *in vitro* phenotypes of *P. aeruginosa* isolates can be related to disease status in CF patients (50). Patient information and status of *P. aeruginosa* infections is limited for the isolates described here, so a comparison between phenotypes and patient status is not feasible. Interestingly, two isolates, UUH0201 and UUH0202, were obtained five months apart from the same patient, with UUH0201 collected first. The UUH0202 strain was less motile and RpoN protein levels dropped compared to UUH0201, but virulence increased. This supports the concept that *in vitro* phenotypes reflect *P. aeruginosa* infection status in CF patients (50). Further work would be needed to fully elucidate such correlations.

The RpoN* molecular roadblock reduced virulence phenotypes in patient isolates with high or low levels of RpoN. For instance, RpoN protein levels were higher in SCH0057-7 than PAO1-S, and RpoN* reduced flagellar and pili motility, biofilm formation and pathogenesis. In contrast, relative RpoN protein levels were low in SCH0256-1 and SCH0354-1, and yet RpoN* reduced motility. Thus, the roadblock was effective in the presence or absence of the native sigma factor. This confirms our previous findings, which show that RpoN* reduced virulence in a laboratory strain that was deleted for *rpoN* (38). Unfortunately, barriers to transformation precluded evaluating RpoN* in some of the other clinical isolates. However, the strains that were successfully transformed represented much of the diversity across the patient isolates.

The CF patient isolates demonstrated variable pathogenesis in the *C. elegans* paralytic killing model that spans 6 hours. Only one pathogenic isolate, SCH0057-7, was transformable and thus possible to evaluate the effects of RpoN* *in vivo*. This strain and several others were also pathogenic in the 4-day slow killing assay, but this assay was not used to evaluate RpoN* because of difficulty maintaining the plasmid and RpoN* expression. Gentamicin selection and IPTG induction are not durable, we found (38), because the *C. elegans* cuticle is impermeable and the compounds are poorly absorbed in the intestine (51). While it is expected that expressing RpoN* in CF isolates would improve *C. elegans* survival in the slow killing assay, it is not feasible with the current vector. If issues with maintaining the plasmid and expression of the roadblock were resolved, it would be interesting to evaluate RpoN* in this assay using patient isolates.

The molecular roadblock, RpoN*, binds numerous promoters in bacterial genomes, altering the transcriptome. RpoN* expression in *P. aeruginosa* greatly reduced transcription of the *mex* family genes (38), which are involved in multidrug efflux pumps (20). Increased expression of *mex* genes is linked to increased resistance to antibiotics (17). Therefore, we investigated whether RpoN* alters *P. aeruginosa* susceptibility to antibiotics. We employed a clinical laboratory assay for testing bacterial susceptibility or resistance to antibiotics, and found that RpoN* improved antibiotic susceptibility at least two-fold for five different antibiotics, including imipenem. This agrees with work by others that showed RpoN is involved in *P. aeruginosa* tolerance of carbapenems, quinolones, and tobramycin (34–36). Unfortunately, the commercial assay uses predetermined antibiotic concentrations in a 96-well plate, limiting the scope of the molecular roadblock’s effects. Additionally, the *P. aeruginosa* strain used here is sensitive to quinolones and tobramycin, so the effects of RpoN* expression on resistance to these antibiotics was not evaluated. It will be important to test clinical strains that are resistant to quinolones, carbapenems, and tobramycin to determine the effects of RpoN*. Further studies are needed to uncover the full spectrum of RpoN* effects on antibiotic susceptibility.

Multi-drug resistant organisms (MRDOs) are increasing worldwide, even those with resistance to entire classes of antibiotics. Alarmingly, nearly all antibiotics brought to market in the past 30 years are variations on existing drugs (52). Research into alternative strategies to treat bacterial infections is a priority, including compounds to enhance the activity of existing antibiotics or neutralize virulence factors. The molecular roadblock falls into the latter type. RpoN* binds consensus promoters throughout the *P. aeruginosa* genome, affecting the transcription of numerous virulence factors. Due to the many binding sites for RpoN*, it is unlikely antibiotic resistance will develop during treatment. The binding sequence of the RpoN consensus promoter is conserved across gram-negative and gram-positive bacteria (36, 37). We explored the effects of RpoN* on virulence phenotypes of *Pseudomonas putida*, *Burkholderia cepacia*, and *Escherichia coli* (unpublished data), suggesting that RpoN* may reduce virulence in multiple organisms. More studies are needed to identify the spectrum of RpoN* activity and its resistance frequency. Currently, the molecular roadblock is a tool for antimicrobial development and is not a usable drug. However, a small molecule that works in the same cis-acting manner as RpoN* would be an effective, clinically relevant strategy to combat *P. aeruginosa* virulence and antibiotic resistance.

## Materials and Methods

### Bacteria and Nematodes

*P. aeruginosa* clinical isolates were provided by the Seattle Children’s Hospital (SCH strains) and Upstate University Hospital (UUH strains). *P. aeruginosa* PAO1-M was provided by C. Manoil (44), and *P. aeruginosa* PAO1-S and Δ*rpoN* were provided by D. Haas (31). *P. aeruginosa* PA19660 *Xen5* was purchased from PerkinElmer. *E. coli* OP50 was provided by D. Pruyne (SUNY Upstate Medical University). All strains are listed in Table 1. For long-term storage, bacteria were grown overnight in LB broth at 37°C with shaking, and frozen in 10% glycerol at −80°C. *Caenorhabditis elegans* N2 was purchased from the *Caenorhabditis* Genetics Center (University of Minnesota, Minneapolis, MN), and maintained on nematode growth media (NGM) seeded with *E. coli* OP50 at 20°C (53). Populations were synchronized via egg lay and grown to the young adult stage at 20°C (54).

### Plasmids

RpoN* and empty vector plasmids were previously described (38). Plasmids were maintained in *E. coli* INV110 (Invitrogen) with gentamicin selection (30 mg/L). RpoN* expression was induced with isopropyl β-D-1-thiogalactopyranoside (IPTG, 1 mM).

### Transformation

Permissive *P. aeruginosa* patient isolates were transformed by electroporation prior to all experiments, per standard protocol (55). Transformed bacteria were selected on LB agar or BHI agar supplemented with gentamicin (30 mg/L). Individual colonies were picked for each assay.

### Western Blot Analysis

Overnight bacteria cultures were treated with Cell Lytic B Lysis Reagent (Sigma) to generate crude cell lysates. The soluble protein fraction was separated on 10% Mini-PROTEAN^®^ TGX Stain-Free^TM^ protein gels (BioRad), activated for 5 minutes with UV light, imaged and transferred via semi-dry apparatus to a PVDF membrane. Membranes were incubated with primary antibody specific for *E. coli* RNA σ^54^ (1:500, BioLegend) overnight, then with secondary antibody HRP goat anti-mouse (1:10,000, Jackson ImmunoResearch). The chemiluminescent signal was generated with the Pierce SuperSignal West Fempto substrate kit (Thermo Scientific), and detected with ChemiDoc™ MP Imaging System (Bio-Rad Laboratories). Protein bands and total protein per lane were measured with Image Lab (Version 5.2.1; Bio-Rad Laboratories). RpoN bands were then compared to corresponding total detected protein in each lane.

### Phenotyping Assays

Assays to measure swimming, twitching (56), and biofilm formation (57), were conducted according to standard protocols. Transformed *P. aeruginosa* clinical isolates were grown in appropriate media supplemented with gentamicin (30 mg/L), and with or without IPTG (1 mM). Motility assay and microtiter plate biofilm assay were conducted at 37°C for 24h. Images of motility assays were obtained with IVIS-50™ (Perkin Elmer) and colony diameter was measured with Living Image software (Perkin Elmer). Biofilms were stained with 0.1% crystal violet, extracted in 95% ethanol, and absorbance was measured at 550 nm with a μQuant microplate spectrophotometer (BioTek).

### P. aeruginosa – C. elegans infection assays

For the paralytic killing assay, laboratory strains, clinical isolates, or transformed *P. aeruginosa* were spread on Brain Heart Infusion (BHI) agar (Difco) with, when applicable, gentamicin (30 mg/L) and with or without IPTG (1 mM). *E. coli* was spread on BHI agar. All plates were grown overnight at 37°C. Bacteria colonies were swabbed onto BHI agar, supplemented with gentamicin and/or IPTG (1 mM) when applicable, and grown at 37°C for 24 h (44). Adult *C. elegans* were added to plates and the assay was conducted at room temperature, per standard protocol (44). For the slow killing assay, laboratory strains or clinical isolates of *P. aeruginosa* were grown overnight in LB broth at 37°C with shaking, and cultures were spread on a modified NGM agar (0.35% bactopeptone, 2% bactoagar) (58). Plates were incubated at 37°C for 24 h, then at room temperature for an additional 24 h. The assay was conducted at 20°C, and worms were scored every 24 h per standard protocol (58).

### Antibiotic Sensitivity Testing

Transformed or mock transformed bacteria were grown overnight in LB broth with gentamicin (30 mg/L) and IPTG (1 mM) or only IPTG (1 mM), respectively. MicroScan Neg MIC 43 panels (Beckman Coulter Inc., Brea, CA) were used. Panels were set up per manufacturer’s protocol (MicroScan Gram Negative Procedure Manual, version 09/2016) using the RENOX system (Beckman Coulter Inc., Brea, CA) with a final well concentration of 3-7×10^5^ CFU/mL. The following modifications were made to the manufacturer’s protocol: LB broth supplemented with IPTG (1 mM) and with or without gentamicin (30 mg/L) was used in place of saline for whole panel. Plates were incubated at 35°C for 16-20 h, and read using a MicroScan autoSCAN-4 (Beckman Coulter Inc, Brea, CA). Quality control was performed on the panels per manufacturer’s protocol.

### Statistics

Data were analyzed using Excel and GraphPad Prism with a significance of *p* ≤ 0.05 (Microsoft, Washington; GraphPad Software Inc., California).

### Data Availability

The datasets produced for the current study are included in this manuscript or are available from the corresponding author upon reasonable request.

## Acknowledgements

This work was supported by the NIH (CTN: 2R15GM104880) and by the Hill Collaboration [58482 to JFM and CTN]. The funders had no role in study design, data collection and interpretation, or the decision to submit the work for publication. *C. elegans* strains were provided by the CGC, which is funded by NIH Office of Research Infrastructure Programs (P40 OD010440). We would like to thank the Clinical Microbiology Lab at the Syracuse VA Medical Center in Syracuse, NY for supplying the antibiotic susceptibility testing panels and for use of their facility for reading the results. We would also like to thank Dr. Ran Anbar and Donna Linder at the Golisano Center at Upstate University Hospital and Marcella Blackledge and Dr. Rafael Hernandez at the Seattle Children’s Hospital for supplying the *P. aeruginosa* CF patient isolates (Seattle Children’s’ Hospital CF patient isolates obtained under NIH P30 DK089507).

## Competing Financial Interests

The authors declare no competing financial interests.

## Author Contributions

M.G.L. wrote the manuscript. C.T.N. and J.F.M. conceived the study. M.G.L. and J.L.V. conducted the experiments. M.G.L. generated the figures. All authors reviewed the manuscript.

